# Stimulus transformation into motor action: dynamic graph analysis reveals a posterior-to-anterior shift in brain network communication of older subjects

**DOI:** 10.1101/2020.02.26.966325

**Authors:** Nils Rosjat, Bin A. Wang, Liqing Liu, Gereon R. Fink, Silvia Daun

## Abstract

Cognitive performance slows down with increasing age. This includes cognitive processes that are essential for the performance of a motor act, such as the slowing down in response to an external stimulus. The objective of this study was to identify aging-associated functional changes in the brain networks that are involved in the transformation of external stimuli into motor action. To investigate this topic, we employed dynamic graphs based on phase-locking of Electroencephalography signals recorded from healthy younger and older subjects while performing a simple visually-cued finger-tapping task. The network analysis yielded specific age-related network structures varying in time in the low frequencies (2-7 Hz), which are closely connected to stimulus processing, movement initiation and execution in both age groups. The networks in older subjects, however, contained several additional, particularly interhemispheric, connections and showed an overall increased coupling density. Cluster analyses revealed reduced variability of the subnetworks in older subjects, particularly during movement preparation. In younger subjects, occipital, parietal, sensorimotor and central regions were - temporally arranged in this order - heavily involved in hub nodes. Whereas in older subjects, a hub in frontal regions preceded the noticeably delayed occurrence of sensorimotor hubs, indicating different neural information processing in older subjects.

All observed changes in brain network organization, which are based on neural synchronization in the low frequencies, provide a possible neural mechanism underlying previous fMRI data, which report an overactivation, especially in the prefrontal and pre-motor areas, associated with a loss of hemispheric lateralization in older subjects.

## Introduction

Aging leads to a slowing down of cognitive abilities. This includes cognitive processes that are crucial for motor performance. For example, increased deficits in the execution and planning of movements, as well as a slowing down in response to external stimuli (Wu & Hallett, 2005; Seidler et al., 2010; Liu et al., 2017). A study by Niessen et al. (Niessen, Fink, Hoffmann, Weiss, & Stahl, 2017) recently showed that older subjects have deficits primarily in the accuracy of processing visual stimuli into motor actions and less in the pure execution of the motor act itself. Even though several studies investigated the effects of non-pathological aging on human motor performance, to date, our understanding of the neurobiological underpinnings of aging-asociated functional changes in the brain networks that are involved in the transformation of external stimuli into movements remains sketchy.

Several fMRI studies reported an association between aging and changes in movement-related neural activity. In particular, a generally enhanced activation was reported, especially in pre-frontal and pre-motor areas (Sailer, Dichgans, & Gerloff, 2000; Heuninckx, Wenderoth, Debaere, Peeters, & Swinnen, 2005; Ward, Swayne, & Newton, 2008). This effect is often referred to as the so-called PASA (posterior to anterior shift in aging) phenomenon (Dennis & Cabeza, 2011). The HAROLD (hemispheric asymmetry reduction in older subjects) model proposes another effect often reported by fMRI studies, namely a reduced hemispheric asymmetry during cognitive processing, which is thought to be rather a general aging-associated than task-specific phenomenon (for a summary see (Cabeza, 2002)). It is assumed that these effects (PASA and HAROLD) reflect either a compensatory mechanism, i.e., a recruitment of additional brain areas to support weakened functionality in core brain regions (Cabeza et al., 1997; Nolde, Johnson, & Raye, 1998; Reuter-Lorenz et al., 2000), or difficulties in recruiting specialized task-specific subnetworks in older subjects (Mitrushina & Satz, 1991; Babcock, Laguna, & Roesch, 1997; Cabeza, 2002). However, fMRI suffers from low temporal resolution. Therefore fMRI-based analyses of functional connectivity between brain regions provide only limited insight into the underlying neural interactions. In this study we intended to shed light on the neural mechanisms underlying HAROLD and PASA using Electroencephalography (EEG)-based functional connectivity analyses taking advantage of EEGs’ high temporal resolution.

One hypothesis states that such changes in healthy aging result from changes in inter-regional neural synchronization since this is a crucial mechanism underlying motor and cognitive tasks (Fell et al., 2004; Popovych et al., 2016; Palva & Palva, 2012). Several studies using data from EEG/MEG/single-unit/ECog recordings while subjects performed a variety of different motor tasks showed that remote neural populations synchronize over a short-time period (Singer, 1999, 2004; Uhlhaas, Roux, Rodriguez, Rotarska-Jagiela, & Singer, 2010; van Wijk, Beek, & Daffertshofer, 2012), suggesting that coordinated timing constitutes a fundamental principle involved in motor and cognitive processing (Uhlhaas et al., 2010; Fries, 2005, 2015; Baker, Olivier, & Lemon, 1997; Baker, Spinks, Jackson, & Lemon, 2001). Furthermore, we could show in a previous study (Rosjat et al., 2018) that during *voluntary*, i.e. internally-triggered movements, motor-related networks built on inter-regional neural synchronization in the lower frequencies in younger as well as older subjects. In addition, the results exhibited an increase in interhemispheric connectivity in older subjects compatible with the HAROLD model. However, due to the restriction of the phase-locking analysis to EEG electrodes above the motor cortex, we were unable to find evidence for the PASA phenomenon.

Accordingly, we here aimed at investigating how changes related to healthy aging affect the dynamic organization of brain networks with a particular interest in the role of occipital, parietal and prefrontal and premotor regions besides the classical motor-related ones. In addition, we focused on visually-cued movements for the following reasons. First, we expected a more distinct effect of PASA in this kind of task, since a raise in cognitive load is known to increase the PASA phenomenon (e.g. (Ansado, Monchi, Ennabil, Faure, & Joanette, 2012)). Second, we were interested in how visual information is transformed towards motor execution in such visually-cued movements. For the investigation of this aspect it is essential to obtain information about the *timecourse* of the network structure. Thus, we decided to focus our analysis on *dynamic graphs*. A dynamic graph (also dynamic network) is a composition of temporally successive connectivity networks (Sizemore & Bassett, 2018; Holme & Saramäki, 2012). In contrast to static networks, they take into account the temporal development of the connectivity. They, therefore, serve to provide a more precise temporal account of the different coupling structures of the respective age groups, especially about the influence of aging on the routing of neural information during the different movement phases. Third, we wanted to analyze whether the neural evidence for the HAROLD model found in voluntary, i.e. self-initiated movements (Rosjat et al., 2018) could be replicated in visually-cued movements, i.e. whether the neural underpinnings are independent of the way the movement was initiated.

We investigated inter-channel connectivity using the relative phase-locking value (rPLV) (Lachaux, Rodriguez, Martinerie, & Varela, 1999) applied to EEG data. We chose to use the rPLV as connectivity measure as it is, in contrast to other methods like dynamic causal modeling (DCM) or Granger causality, capable of measuring fast transient synchronization. The high temporal resolution of this connectivity measure is crucial for investigating the underlying dynamic neural networks in our study. EEG data were recorded from younger and older healthy participants who performed a simple visually-cued finger-tapping task and dynamic networks based on these phase relationships were created.

## Materials and Methods

### Participants

In this study, we included EEG data of a group of twenty-one younger healthy subjects (YS; 10F/11M, age: 22-35 years) and of a group of thirty-one older healthy subjects (OS; 15F/16M, age: 60-78 years) first presented in two earlier studies (Popovych et al., 2016; Liu et al., 2017). All participants were right-handed according to the Edinburgh Handedness Inventory (Oldfield, 1971) and had normal or corrected to normal color vision. The older participants had no history of psychiatric or neurological disease (as assessed by the Trail making test (TMT A: 41.16 ± 16.27 s, TMT B: 78.59 ± 28.02 s) (Spreen & Strauss, 1998), the Mini-mental-state-test (28.97 ± 1.29) (Folstein, Folstein, & McHugh, 1975), the Clock-drawing test (Agrell & Dehlin, 1998), and the Beck depression inventory (4.58 ± 3.21) (Beck, Ward, Mendelson, Mock, & Erbaugh, 1961). All participants had given their written informed consent prior to the study. The local ethics committee of the Faculty of Medicine at the University of Cologne had approved both studies.

### Experimental Design

We recorded EEG data while both groups performed a simple finger-tapping task. The experimental paradigm reported here consisted of two conditions, a visually-cued condition with intention to act, i.e. a visually-cued motor condition (Fig. 1 (A)) and a visually-cued condition without intention to act, i.e. a vision-only condition (Fig. 1 (B)). In the motor condition (visually-cued tapping, Fig. 1 (A)) the subjects were presented with a right- or left-pointing arrow (2°wide and 1.2°high, expressed as visual angles) for 200 ms on a screen with inter-stimulus intervals of varying length ≥ 4 s. The participants had to press a button with their left or right index finger, corresponding to the direction of the arrow, as fast as possible. In the second condition (vision-only, Fig. 1 (B)) the same stimuli as in the visually-cued tapping condition were presented. However, this time the participants were carefully instructed to pay attention to the arrows but to refrain from performing or imagining to perform the button press. Thus, no motor action was performed in this condition (for full details see (Popovych et al., 2016; Liu et al., 2017)). During the experiment, a third, self-initiated tapping condition (not reported here) was recorded. In this condition, the participants were free to choose when to move which hand without any external stimuli. The conditions were presented in pseudorandomized blocks of one minute duration each with a break of 17-20 s between consecutive blocks (for details see (Wang et al., 2017)). The overall duration of the experiment including breaks was 70 min.

**Fig 1.**
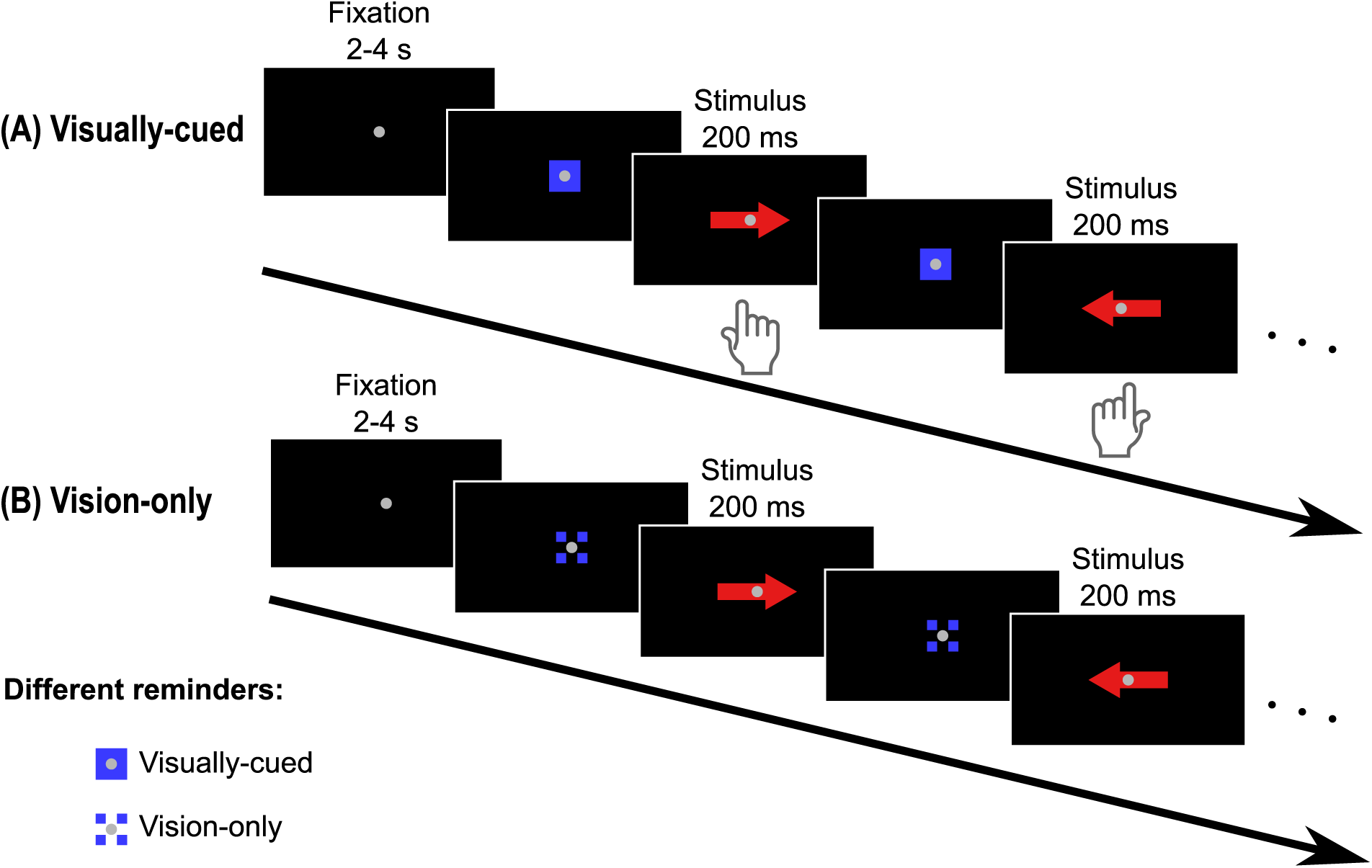
Experimental paradigm. Organizational setup of the experiment. (A) Visually-cued finger movements and (B) vision-only condition. Each condition is represented by a unique reminder (e.g. blue square in the visually-cued tapping condition). The arrows indicate the hand (left or right) to be used (adapted from Fig. 1 in (Wang et al., 2017)).

Our analyses were restricted to data acquired from the visually-cued and vision-only condition (Fig. 1 (A) and (B)) and the conditions were analyzed locked to the onset of the visual stimulus. We used the same cohort as in (Rosjat et al., 2018) for the following reason. We wanted to check whether the type of movement initiation (internal versus external) has an influence on the observed coupling structure. Therefore, we decided to leave the cohort constant to ensure that any differences, e.g. in phase-locking are due to changes in the experimental task.

### EEG recording and preprocessing

While the subjects performed the task, we continuously collected EEG data from 64 active Ag/AgCl electrodes (Brain Products GmbH, Munich, Germany), placed according to the international 10-20 system. The reference electrode was placed at the left earlobe. Bipolar horizontal and left vertical electrooculograms (EOG) were recorded with three of the 64 scalp electrodes (previously located at FT9, FT10, and TP10 in the 10-20 system). These were placed at the bilateral outer canthi and under the left eye to monitor eye movements and blinking. Before the experiment, it was ensured that the impedances of all electrodes were below 15 kΩ. The EEG signals were amplified, bandpass filtered in the frequency range 0.87 − 500 Hz, and digitized at a sampling rate of 2.5 kHz. Index finger movement onsets were detected by acceleration sensors attached to the tip of each index finger. We also used the information from the acceleration signals to monitor the subjects’ behavior, e.g., to rule out errors such as mirror movements. We defined the onset of the finger movement as the instant of time at which the numerical time derivative of the acceleration signal exceeded a predefined threshold.

The acquired raw data went through several offline pre-processing steps. First, we band-pass filtered the data from 0.5 to 48 Hz to remove slow voltage drifts. We next downsampled the data to 200 Hz to reduce the file size and thus the computing time. We then removed artifacts using a semi-automated process in EEGLAB using independent component analysis (ICA) (Makeig, Bell, Jung, & Sejnowski, 1996). Finally, the data were epoched to intervals [−1500, +2500] ms, which centered around stimulus onset.

For our analysis, we excluded trials that contained EEG artifacts or included movements during the pre-stimulus period detected by the accelerometer. After excluding trials based on these criteria, we only considered data from participants who had at least 30 correct trials per hand. Based on these quality considerations, data from three younger and seven older subjects were excluded from further analysis. Thus, EEG data from 18 younger and 24 older subjects were used for further processing.

### Spatial filtering

A crucial pre-processing step for the following connectivity analysis was spatial filtering. In the EEG, volume conduction is a primary concern in determining the relationship of signals. The activity of a single neural generator can influence the signal in several different electrode positions and thus lead to inaccurate phase-locking between these recording sites. To reduce the detection of false-positive connectivity, we applied spatial filtering using surface Laplacians, which demonstrably improves spatial resolution and allows the analysis of electrodes close to the region of interest (Lachaux et al., 1999). Thus, our preprocessed data were re-referenced to a small Laplacian reference to improve the spatial resolution of the signals and thus their suitability for subsequent connectivity analysis (Hjorth, 1975). Electrodes on the outer edge, which do not have four neighbours, were excluded from further analyses, so that a total of 42 electrodes were included in the investigation.

### Relative amplitude changes and phase-locking in oscillations

Following preprocessing, we transformed the epoched and cleaned data to the time-frequency domain using Morlet wavelets (Kronland-Martinet, Morlet, & Grossmann, 1987) in a *δ* − *β* frequency range (2-30 Hz) with a step size of 1 Hz (5 cycles). This time-frequency decomposition was performed with the Statistical Parametric Mapping toolbox (SPM12, Wellcome Trust Centre for Neuroimaging, London UK) implemented in MatLab R2018b (The MathWorks Inc., Massachusetts, USA). We then analyzed the amplitude and phase information from the time-frequency decomposition using customized MATLAB scripts.

We obtained the temporal evolution of the amplitude A(*f, t*) and phase *ϕ*(*f, t*) by applying the complex Morlet wavelet transformation for each frequency separately to our data. We analyzed the relative amplitude changes for the *δ* − *θ* (2-7 Hz), *α* (8-12 Hz) and *β* (13-30 Hz) frequency ranges separately for the visually-cued motor task and for the contrast between the visually-cued motor task and the vision-only task (i.e. visual stimulation without intention to move).

We quantified the communication of two different brain regions, i.e., acquisition sites, by synchronization determined by the single-frequency phase-locking value (sPLV; adapting the phase-locking value defined in (Lachaux et al., 1999)). For a pair of channels m and n, sPLV is defined as:

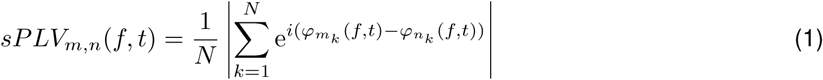

here 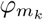 denotes the phase of the EEG at channel m in the k-th trial. N is the total number of trials, and i is the imaginary unit. While minimum sPLV = 0 represents a random distribution of phase differences over all trials without any coherence, a maximal sPLV = 1 occurs only in the case of perfect inter-trial phase locking of the phase differences between the EEG signals at the two channels m and n over all trials.

Since we were interested in an event-related measure that represented the synchronization increases and decreases during stimulus processing, movement preparation and execution, we normalized the sPLV of the electrophysiological recordings at each pair of channels with respect to its baseline value and calculated its relative change over the entire epoch. We refer to these normalized sPLVs as *relative phase-locking values* (rPLVs):

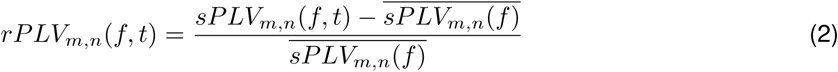

here 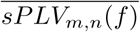 denotes the sPLV of the baseline interval for each frequency, i.e. the mean sPLV at frequency f in the baseline interval, which was defined from [−1300, −500] ms excluding the first 200 ms of the original interval as it contained edge artifacts of the Morlet transformation.

To analyze inter-regional synchronization, we considered four major frequency bands: *δ* (2-3 Hz), *θ* (4-7 Hz), *α* (8-12 Hz) and *β* (13-30 Hz). Instead of first filtering the signal to a specific frequency band, we calculated the rPLV separately for each frequency with a resolution of 1 Hz before averaging these values over the frequencies of the frequency band. In this way, we ensured a true 1:1 frequency coupling that maximizes the contribution of each frequency in the band. The steps described above were performed for both the tapping and the vision-only control condition. The resulting mean rPLVs (averaged over a frequency band of interest) were then contrasted by subtracting the vision-only control condition from the visually-cued condition, resulting in positive values when phase-locking is stronger in visually-cued and negative values for stronger phase-locking in the control condition. (cf. Fig. 2). The contrasted mean rPLVs then underwent statistical testing (described below) to define the edges of the phase-locking network. The resulting networks were analyzed using graph theoretical metrics as implemented in the Brain Connectivity Toolbox (Rubinov & Sporns, 2010).

**Fig 2.**
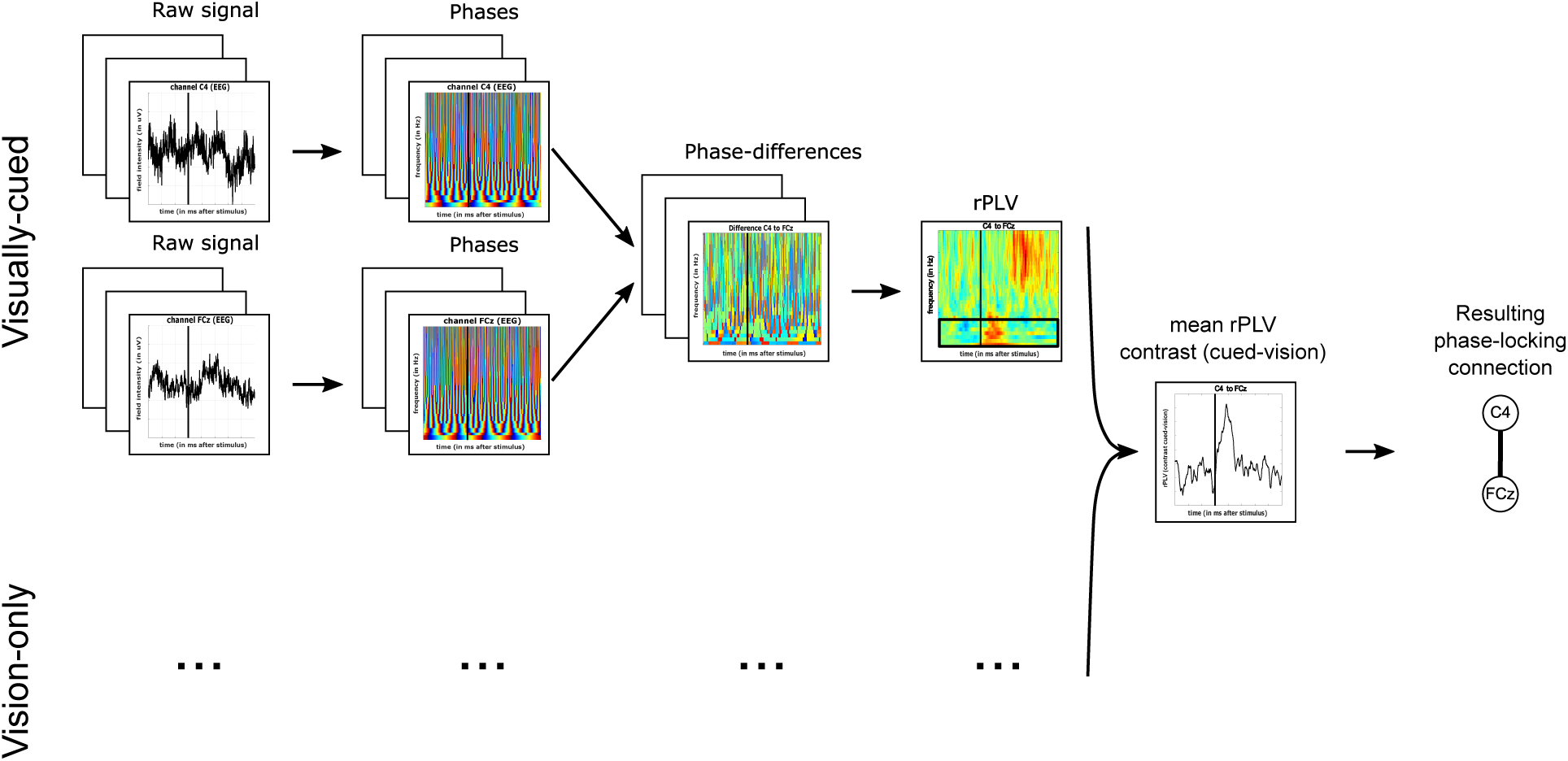
Phase-locking analysis. Schematic illustration of the procedure to construct corresponding phase-locking networks from contrasted, i.e., visually-cued - vision-only, rPLVs (adapted from Fig. 1 in (Rosjat et al., 2018)). The black vertical lines represent movement onset.

By implementing the contrast condition we reduced the pure sensory effect of the visual stimulation, which would otherwise have to led to an all-to-all connectivity in the networks.

### Statistical analysis

We applied pointwise t-tests with a significance level of *p <* 0.05, corrected for multiple comparisons (false discovery rate, FDR, *q* = 0.05) (Benjamini & Hochberg, 1995), to test whether the mean relative amplitude changes (averaged over frequency band scaled logarithmically relative to pre-stimulus baseline) between YS and OS differed significantly in the intervals [0, 1000] ms (*δ* − *θ*) and [0, 1500] ms (*α*). To test whether the difference between the mean rPLV (averaged over the frequency band) and the baseline (averaged over the frequency band) was statistically significant, we compared the mean rPLV obtained for each pair of electrodes at each time point in the interval [0, 1000] ms with its baseline value. To this end, we used a pointwise t-test with a significance level of *p <* 0.05, corrected for multiple comparisons (false discovery rate, FDR, *q* = 0.05) (Benjamini & Hochberg, 1995). The corrections were performed concerning the number of time points, age groups, and electrodes. The baseline was constructed by generating normally-distributed random values that had the same mean and standard deviation as the EEG at any time point of the baseline interval [−1300, −500] ms.

For the construction of the rPLV networks, we defined a *connection*, i.e., an edge of the network, between two given nodes to exist at a given time point t if the mean rPLV between these two nodes was significantly increased with respect to its baseline value. Based on our previous results (Rosjat et al., 2018; Popovych et al., 2016), which showed a symmetrical behavior of phase-locking, and to reduce the number of statistical comparisons, the rPLVs during left- and right-hand finger movements were collapsed together after flipping the ones for the right-hand movements. Electrodes were defined as being either ipsilateral or contralateral to the moved hand. For convenience, results are presented resembling left-hand movements, i.e., electrodes over the left hemisphere are assumed to be ipsilateral while electrodes over the right hemisphere are assumed to be contralateral to the performed movement.

### Setup of dynamic networks

The statistical analysis above defines a binary undirected graph *G*_*t*_ = (*V, E*_*t*_), where the graph G is defined by a set of vertices *V* and edges *E*_*t*_ : *V* × *V* → R, for each point in time t for younger and older participants. The dynamic graph, also called dynamic network, is defined as the ordered set of graphs *G*(*t*) = {*G*_*t*_|*t* ∈ [1, *…, T*]}. It can also more efficiently be represented by a list of triples (*i, j, t*), defining the contacts of two nodes i and j and the time point t at which they occur (Sizemore & Bassett, 2018). In contrast to static graphs, dynamic graphs account for time-varying connectivity patterns that might reflect the formation of subnetworks due to task performance.

### Aggregated networks

Besides the dynamic networks, we investigated the *aggregated graphs* of the dynamic networks, which are graphs that account for the number of occurrences of each edge over all graphs, i.e., edges that appear more often have a higher weight while rarely present edges have a smaller weight.

### Community structure

We performed a cluster analysis using the Louvain community algorithm (clustering parameter *q* = 0.9), which uses a greedy optimization method to minimize the ratio of the number of edges inside communities to edges outside communities, for each time step and the aggregated graph for both groups of participants to retrieve information on closely connected subnetworks (Blondel, Guillaume, Lambiotte, & Lefebvre, 2008; Reichardt & Bornholdt, 2006; Ronhovde & Nussinov, 2009; Sun, Danila, Josić, & Bassler, 2009). To reduce the effect of randomly assigned cluster labels between time points, we always used the previous cluster as the initial condition for the following community detection. Additionally, we post-hoc assigned a label for each community, minimizing the number of label switches between time points, which prevents the same cluster from being assigned different cluster labels at consecutive time points. Furthermore, based on these clusters we tested the similarity of clusters between age groups by computing the variation of information (VI) (Meilă, 2007)

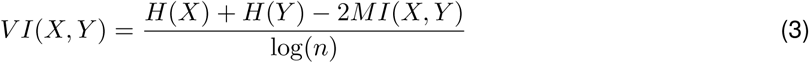

where H is entropy, MI is mutual information, and n is the number of nodes. This metric ranges from 0, i.e., identical clustering, to 1 (as it is normalized by log(*n*)), i.e. maximally distinct clustering. To account for actual age effects, we compared *V I*(*X, Y*) (X representing younger subjects and Y representing older subjects) with time-lagged self-distribution distances *V I*(*X*(*t*), *X*(*t* − 1)) and *V I*(*Y* (*t*), *Y* (*t* − 1)) to test whether the partition distance between younger and older subjects is larger than within group partition changes in time.

The flexibility F of the dynamic network is defined by the average over all *node flexibilities f*_*i*_ (Bassett et al., 2011), which are defined as the number of times that a node changed communities, normalized by the maximal number of times this node could have changed communities,

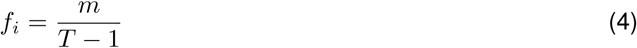

with m the number of community changes and T the number of time points.

### Network information flow

We analyzed the hub nodes of the networks. Those nodes play a crucial role in the network, as they serve as the connection between different subnetworks. For the analysis of hub nodes, we calculated the *betweenness centrality* for each time point, which is defined as the fraction of shortest paths that pass through each node (Brandes, 2001; Kintali, 2008). We primarily focused on the first occurrence of each hub node in the network as well as on their timecourses. This gives an indication of how neural information is transmitted from stimulus processing to motor action.

## Results

### Behavioral results

We first tested whether there was a significant difference between older subjects (OS) and younger subjects (YS) in reaction time (RT), defined as the time that elapsed from stimulus presentation until movement onset, and accuracy rate, i.e., the percentage of correct responses among all trials. The reaction time was significantly longer in older subjects than in younger subjects (see Tab. 1, first row). The mean response accuracy in the visually-cued condition for older subjects and for younger subjects was greater than 90%, consistent with the instructions of the task. However, as expected, the accuracy was lower for the older participants (Tab. 1, second row). We further tested the movement times, i.e. the time from movement onset (determined using the accelerometer) to the actual button press of younger vs. older subjects for significant differences and found that the finger movements of older subjects were significantly (however on average only 6 ms) slower than those of the younger ones (Tab. 1, third row).

In summary, and consistent with the current literature, we found an age-related deceleration in response to the external stimuli and a reduced accuracy in task performance. However, the performance of the motor act itself was only slightly impaired.

### Relative amplitude changes in oscillations

We next investigated the changes in amplitudes in response to the visual stimulus to check whether the data reveal known basic motor-related effects (ERD/ERS) and to clarify whether phase-locking effects could possibly be due to changes in amplitude in the same frequency range. During processing of visual stimulation and execution of movement, both age groups show a relative increase in amplitude in *δ* − *θ* frequencies (2-7 Hz), both in the visually-cued motor and in the contrast condition. Consequently, there are no significant differences in relative amplitude between the two age groups (Fig. 3 A, third column). In the *α* frequency range (8-12 Hz), event-related desynchronization (ERD) can be observed in both groups in both the visually-cued and the contrast condition during motor execution, i.e. around 500 ms after stimulus. In the visually-cued condition, OS show a significantly increased ERD (i.e. greater event-related spectral power decrease) in all examined electrodes compared to YS (2.1202 *<* t’s *<* 5.3578, p’s *<* 0.05 (FDR corrected), df = 40, 0.4202 *<* sd’s *<* 3.2969) (Fig. 3 B top row). Note, that the ERD in OS also lasts longer than in YS, reaching well into the post-movement phase in the central, motor and parietal electrodes (up to 1300 ms, orange bar, 2nd column, Fig. 3B). In the contrast condition, however, the differences in ERD are much less pronounced (all p’s *>* 0.05, apart from few p’s *<* 0.05) (Fig. 3 B bottom row). In *β* frequencies (13-30 Hz), the ERD is also visible in most electrodes during movement. Additionally, especially in YS, we observe an event-related synchronization (ERS) after the execution of the movement (approximately 1000 ms after stimulus). The ERS is significantly stronger in YS than in OS in both, the visually-cued (−3.4455 *<* t’s *<* 6.4240, p’s *<* 0.05 (FDR corrected), df = 40, 0.2788 *<* sd’s *<* 1.4721) and the contrast condition (−3.8928 *<* t’s *<* 5.2834, p’s *<* 0.05 (FDR corrected), df = 40, 0.4394 *<* sd’s *<* 1.4033) (Fig. 3 C, bottom row). In summary, we observe age-related differences in ERD and ERS in the *α* and *β* frequencies and no age-related differences in the *δ* − *θ* frequencies consistent with the current literature.

**Fig 3.**
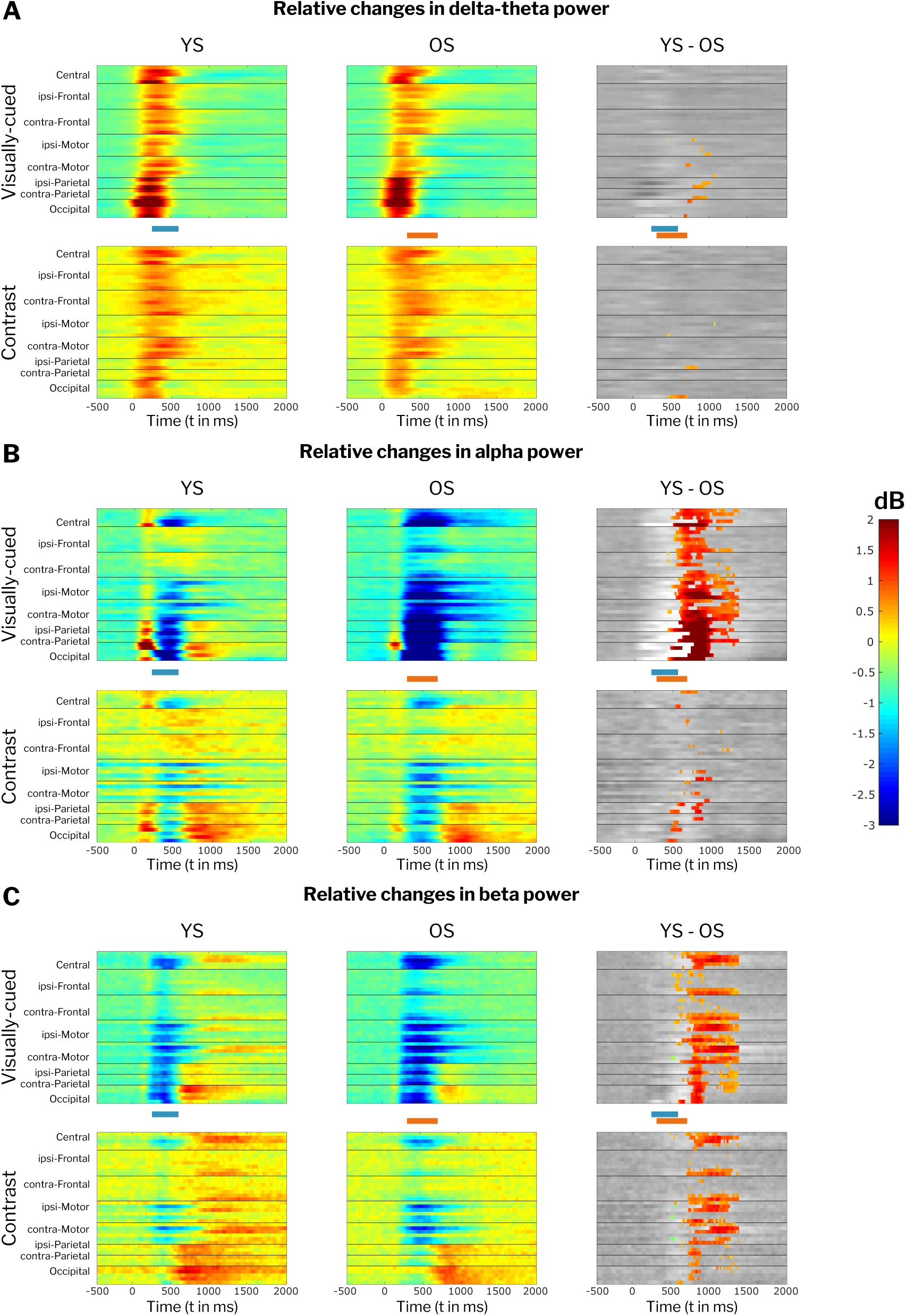
Event-related amplitude changes. Amplitude differences on a logarithmic scale in *δ* − *θ* (A), *α* (B) and *β* frequencies (C) in visually-cued (top rows) and contrast conditions (lower rows) for YS (left column), OS (middle column) and differences YS-OS (right column) for all 42 considered electrodes. Non significant amplitude differences between YS and OS are greyed out in the right column. *t* = 0 represents the onset of the visual stimulus. Blue and orange bars depict time-intervals of motor responses for YS and OS, respectively.

**Table 1.** Summary of statistical results.

|  | mean YS | std YS | mean OS | std OS | t-value | df | p-value |
| --- | --- | --- | --- | --- | --- | --- | --- |
| Reaction time | 426 ms | 77 ms | 497 ms | 84 ms | 3.9442 | 82 | 0.00017 |
| Accuracy | 97.7 % | 2.22 % | 93.13 % | 5.69 % | -3.2329 | 40 | 0.0025 |
| Movement time | 81.9 ms | 8.6 ms | 88.3 ms | 8.3 ms | -3.4307 | 82 | 0.00095 |
| Interhemispheric connections | 3.6 | 3.6643 | 8.55 | 8.5112 | -11.159 | 198 | < 0.0001 |
| Node flexibility (200–300 ms) | 0.3874 | 0.1526 | 0.2043 | 0.1253 | 5.6589 | 40 | < 0.0001 |
| Node flexibility (1–700 ms) | 0.352 | 0.0784 | 0.3094 | 0.0587 | 2.7318 | 40 | 0.0092 |
| Variance of information (YS) | 0.3271 | 0.1425 | 0.5426 | 0.1595 | 29.8243 | 198 | < 0.0001 |
| Variance of information (OS) | 0.338 | 0.1168 | 0.5426 | 0.1595 | 24.9761 | 198 | < 0.0001 |
| Variance of information (YS vs OS) | 0.4321 | 0.0655 | 0.2548 | 0.0593 | 9.6527 | 19 | < 0.0001 |

### Phase synchronization

We found an increased phase-locking in the *δ* − *θ* frequency band (2-7 Hz) during stimulus processing and movement preparation and execution in both age groups and both conditions (Fig. 4). In the visually-cued condition, a significant increase in rPLV was present throughout all electrode pairs (Fig. 4 top row) in both age groups (Ys: −12.3739 *<* ts *<* 5.6742, p *<* 0.05 (FDR corrected), df = 17, 0.0709 *<* sd *<* 2.1193; OS: −11.0582 *<* ts *<* 4.6396, p *<* 0.05 (FDR corrected), df = 23, 0.0942 *<* sd *<* 1.7203), whereas contrasting highlighted more specific connections (Fig. 4 bottom row) (Ys: −8.8947 *<* ts *<* 5.8060, p *<* 0.05 (FDR corrected), df = 17, 0.0987 *<* sd *<* 1.3434; OS: −8.2287 *<* ts *<* 5.8060, p *<* 0.05 (FDR corrected), df = 23, 0.1148 *<* sd *<* 1.1383). In addition, there were increased rPLVs in the *β* frequency band (13-30 Hz) in the post-movement period (approx. 500 ms after stimulus presentation). However, this phase-locking effect in the *β* frequency band could only be observed in few electrode pairs and therefore did not survive the FDR correction. No significantly increased rPLVs were observed in the *α* frequencies. We hence focused our further dynamic graph analysis on the *δ* − *θ* frequencies and on the contrast condition in which the pure sensory effect of the visual stimulation was reduced.

**Fig 4.**
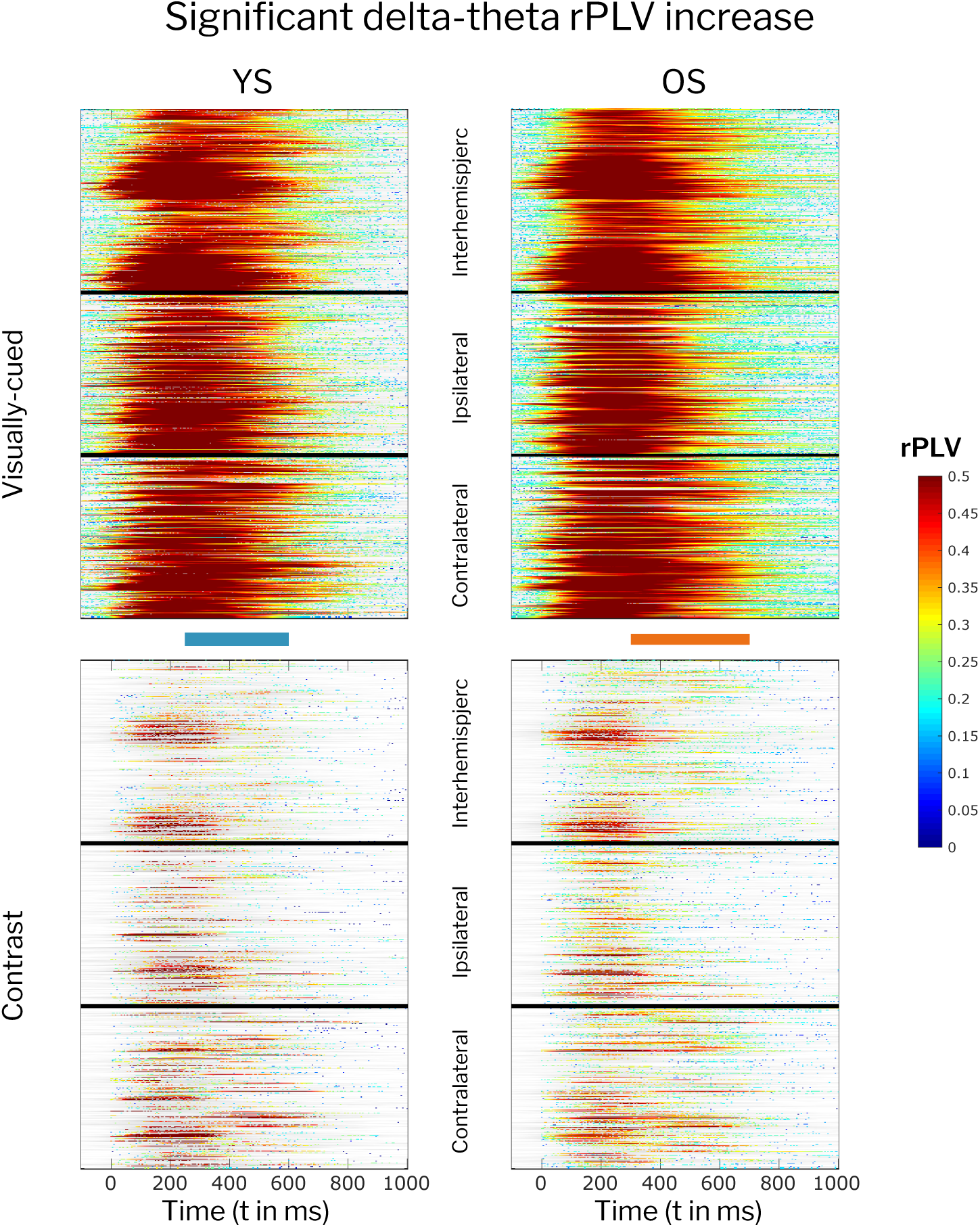
Event-related rPLV changes. Significant increase in phase-locking in the *δ* − *θ* frequency band in the visually-cued condition (top) and in the contrast condition (bottom) for YS (left) and OS (right) for 819 electrode pairs. rPLV values are color coded from 0 (no change) to 0.5 (50% increase). Non significant changes are greyed out. *t* = 0 represents the onset of the visual stimulus. Blue and orange bars depict time-intervals of motor responses for YS and OS, respectively.

### Dynamic networks and graph connection density

Representative graphs resulting from the dynamic network analysis are shown in Fig. 5 (for a video displaying the full timecourse of the dynamic graphs see Supp. V1). The figure displays network connectivity which can roughly be assigned to stimulus processing (150 ms), movement preparation (250 ms), movement execution (350 & 450 ms) and the post movement phase (550 ms). The younger subjects displayed the densest networks in the stimulus processing and movement preparation phase, the older subjects additionally in the early phase of movement execution. The connections in YS networks clustered around occipital regions and central motor-related regions, while OS networks showed a dense connectivity spread over the whole brain. Both group of subjects associated graphs expressed pronounced connectivity between ipsilateral (blue) and contralateral (green) sensorimotor nodes in the movement execution and post movement phase. However, a paired t-test revealed that the older subjects’ networks additionally included an increased number of interhemispheric connections (orange) which were already present in the movement preparation phase (Tab. 1, fourth row). Overall, older subjects’ networks displayed an increase in connectivity, leading to a denser graph.

**Fig 5.**
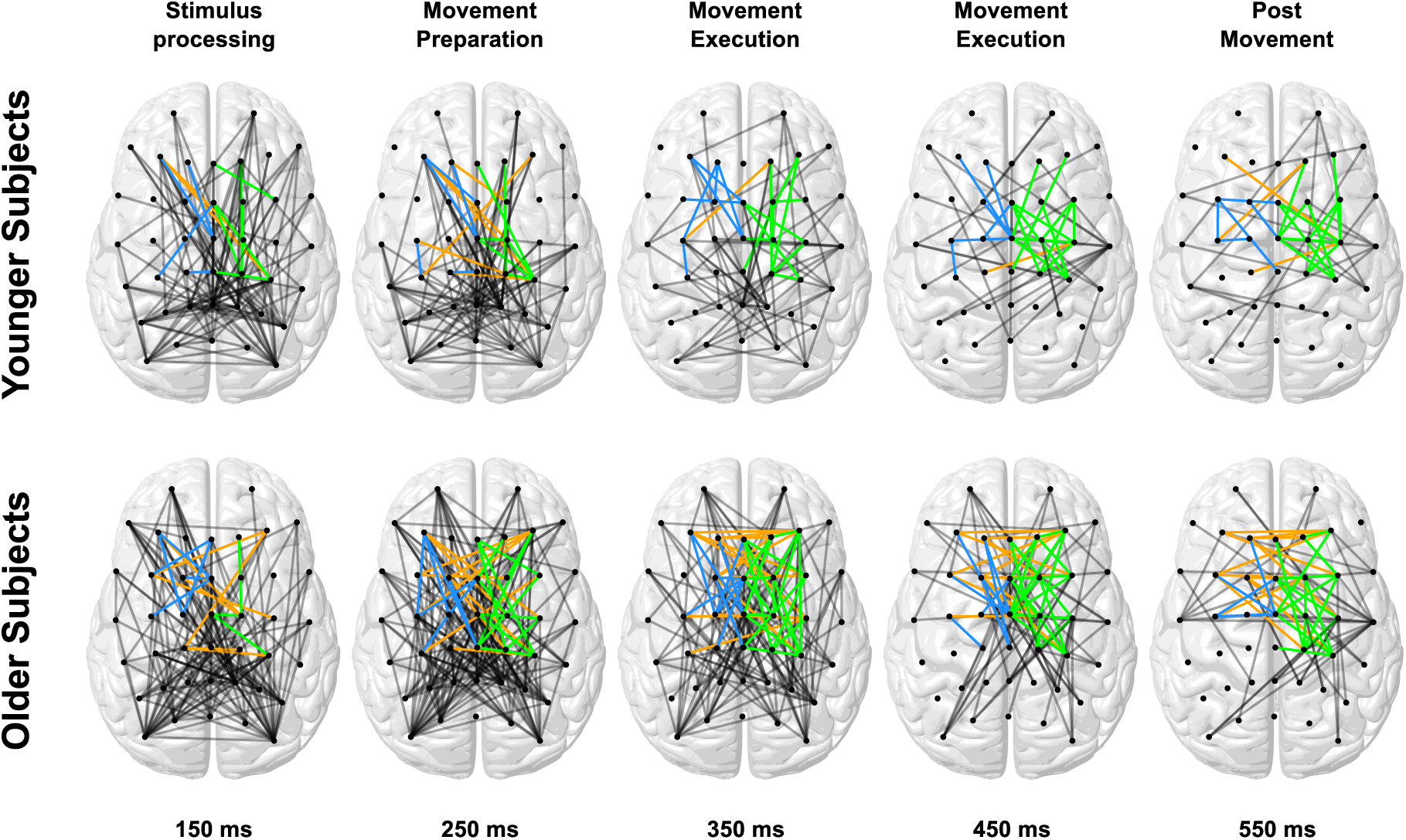
Dynamic graph snapshots in the *δ* − *θ* frequency range. Exemplary graph snapshots for YS (top) and OS (bottom) for five different time points representative for stimulus processing, movement preparation and execution and post movement. Edges related to the core motor network are color coded in blue (ipsilateral), green (contralateral) and orange (interhemispheric). Color codes are chosen as in (Rosjat et al., 2018) for better comparison.

We quantified this by analyzing the average node degree over time for YS and OS (Fig. 6). A paired t-test showed that the average node degree was significantly increased in OS compared to YS in whole network connectivity, i.e., taking all nodes of the network into account (t’s*<* −2.9027, p’s *<* 0.05 (FDR-corrected), df = 60, 0.3404 ≤ sd’s ≤ 4.7489). The same effect was observed for motor network connectivity, i.e., taking only nodes above the motor-related regions into account (t’s *<* −3.3773, p’s *<* 0.05 (FDR-corrected), df = 19, 1.0954 ≤ sd’s ≤ 4.8284). In both cases, the significant differences were mainly present in the time interval from 200 to 400 ms.

**Fig 6.**
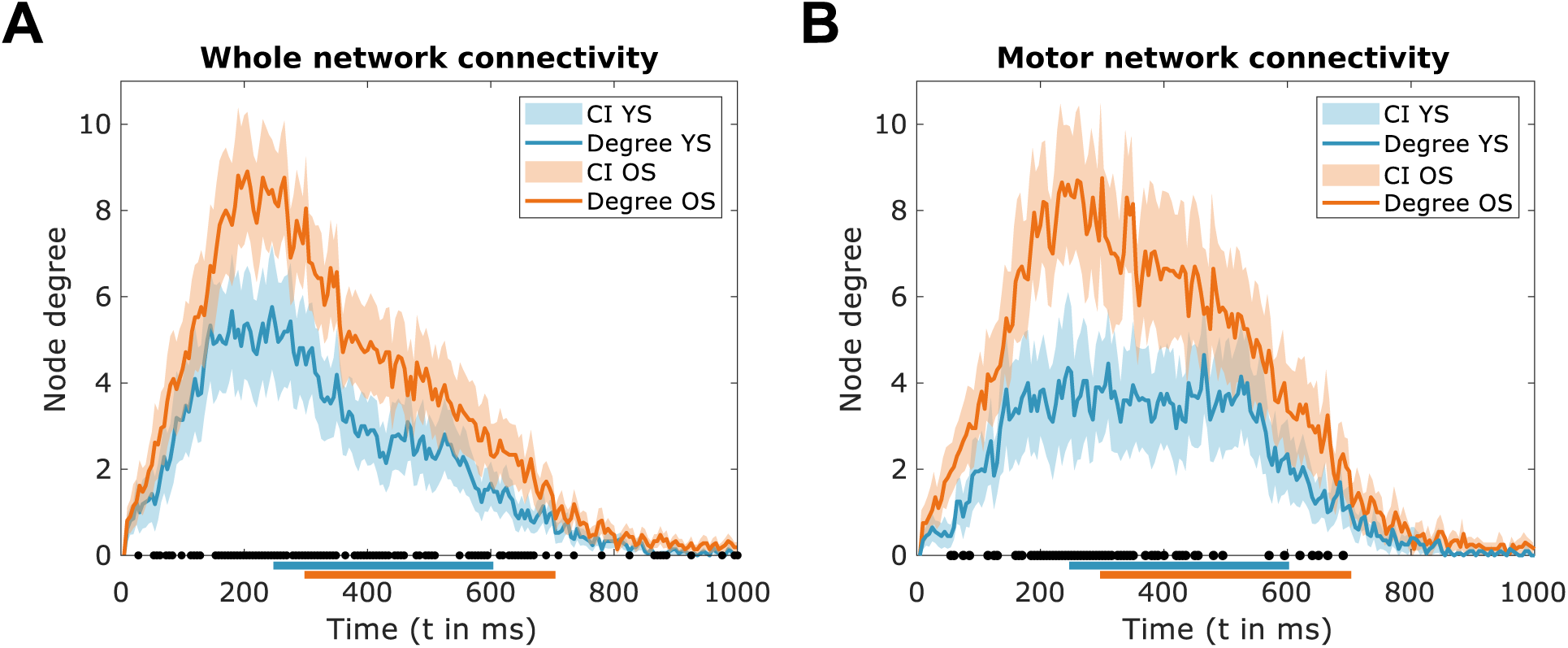
Node degree dynamics. A: Timecourses of the average overall node degrees of YS (blue) and OS (red) surrounded by shaded regions representing the confidence interval. B: Timecourses of the average motor network node degrees of YS (blue) and OS (red) surrounded by shaded regions representing the confidence interval. Movement periods are marked with blue (YS) and red (OS) solid lines at the x-axes. Intervals with significant differences between age groups are marked with black dots.

### Community structure

In the following, we applied the Louvain clustering method (Fig. 7 A, B). During stimulus processing, YS networks were separated in a cluster with a focus on occipital and parietal electrodes (red nodes) and a cluster containing motor, frontal, and central electrodes (light blue nodes). OS networks, on the other hand, were divided into three more widespread clusters with electrodes from almost all eight regions (Fig. 7 B (100 ms)). In the movement preparatory and early movement execution phases (200-300 ms), OS network clusters were reduced to two extended, stable clusters that integrated electrodes from all 8 regions (cf. Fig. 7 B (200 and 300 ms)). The YS networks, on the other hand, broke up into several smaller clusters, also showing no clear regional distinction. A paired t-test showed a significant reduction in node flexibility in OS in the interval (200, 300) ms (Tab. 1, 5th row). Even when the interval was increased to (1-700) ms the node flexibility was significantly reduced in OS (Tab. 1, 6th row). During movement execution the number of connections decreased (cf. Fig. 6 A), which led to a sparser network that for YS could be divided mainly into a contralateral motor - central - ipsilateral frontal community (red) and several smaller communities (Fig. 7 B, 400 ms). In OS, the motor, frontal, and central electrodes were more densely connected, which led to less distinct clusters as in YS.

**Fig 7.**
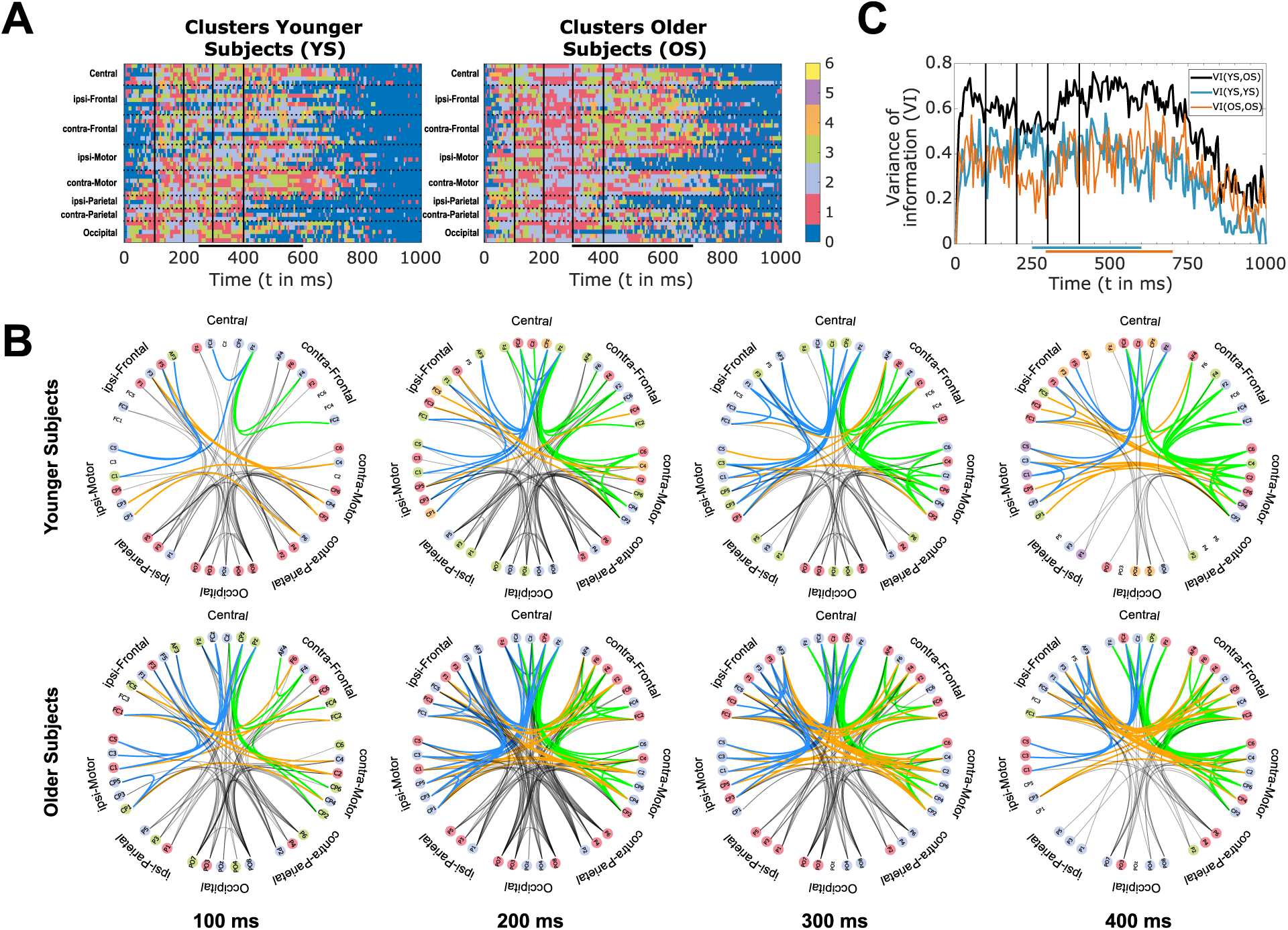
Louvain clustering. A: Channel partitions for YS (left) and OS (right). Clusters are labeled from 1 to 6 (with arbitrary colors). Unconnected channels are displayed in blue (cluster 0). Movement periods are marked with black horizontal lines. B: Circular graphs showing representative clusterings from 100 ms, 200 ms, 300 ms, and 400 ms post stimulus. Nodes are colored corresponding to the clusters in A. Edges related to the core motor network are color-coded in blue (ipsilateral), green (contralateral), and orange (interhemispheric). C: Distribution distance for clusters computed for younger and older subjects (black) compared to self-distribution distances of younger (blue) and older subjects (red). Movement periods are marked with blue (YS) and red (OS) solid lines.

We calculated the variance of information (VI), which serves as a marker of the difference of two clusters of the same nodes, between both groups for each time point (VI(YS,OS)). As a comparison and, in particular, to detect real effects of age, we performed a paired t-test comparing VI(YS,OS) to the VI for YS (VI(YS,YS); Tab. 1, 7th row) and OS (VI(OS,OS); Tab. 1, 8th row) for consecutive time points (Fig. 7 C). With this analysis, we could show that the variance between age groups stays at a high level in the first 700 ms of the epoch (with a value between 0.6-0.8). The variance between age groups was constantly higher compared to the self-distribution distances of YS and OS, except for the time interval [200, 300] ms. In this interval, the self variance of information for OS displayed a drop compared to YS as confirmed by a paired t-test (Tab. 1, 9th row). This drop was related to the less variable networks, i.e., the decreased node flexibility during movement preparation described above. At the same time, YS showed an increased variance of information reflecting a high clustering variability during that time.

### Aggregated networks

In the following, we investigated the aggregated networks, in order compare the results presented here on visually-cued movements, with the already published results of the analyses on self-initiated movements (Rosjat et al., 2018), as these had a much lower time resolution. The resulting networks are presented in Fig. 8. YS and OS showed densely connected nodes above the occipital cortex, which were related to the early time intervals of the dynamic graphs. In YS the motor network showed additionally dense connectivity between nodes above ipsilateral frontal and central regions (blue), diagonal interhemispheric connections between nodes above the ipsilateral frontal and contralateral motor cortex (orange), horizontal interhemispheric connections between nodes above the ipsilateral and contralateral motor cortex (orange), and between nodes above the contralateral motor and central electrodes (green) (Fig. 8 A). In contrast, OS showed a densely connected network between all nodes, but more strikingly an increase in interhemispheric horizontal connectivity between nodes above the frontal, frontocentral, and central regions, i.e., between nodes above the ipsilateral and contralateral frontal and above the ipsilateral and contralateral motor cortex (Fig. 8 B).

**Fig 8.**
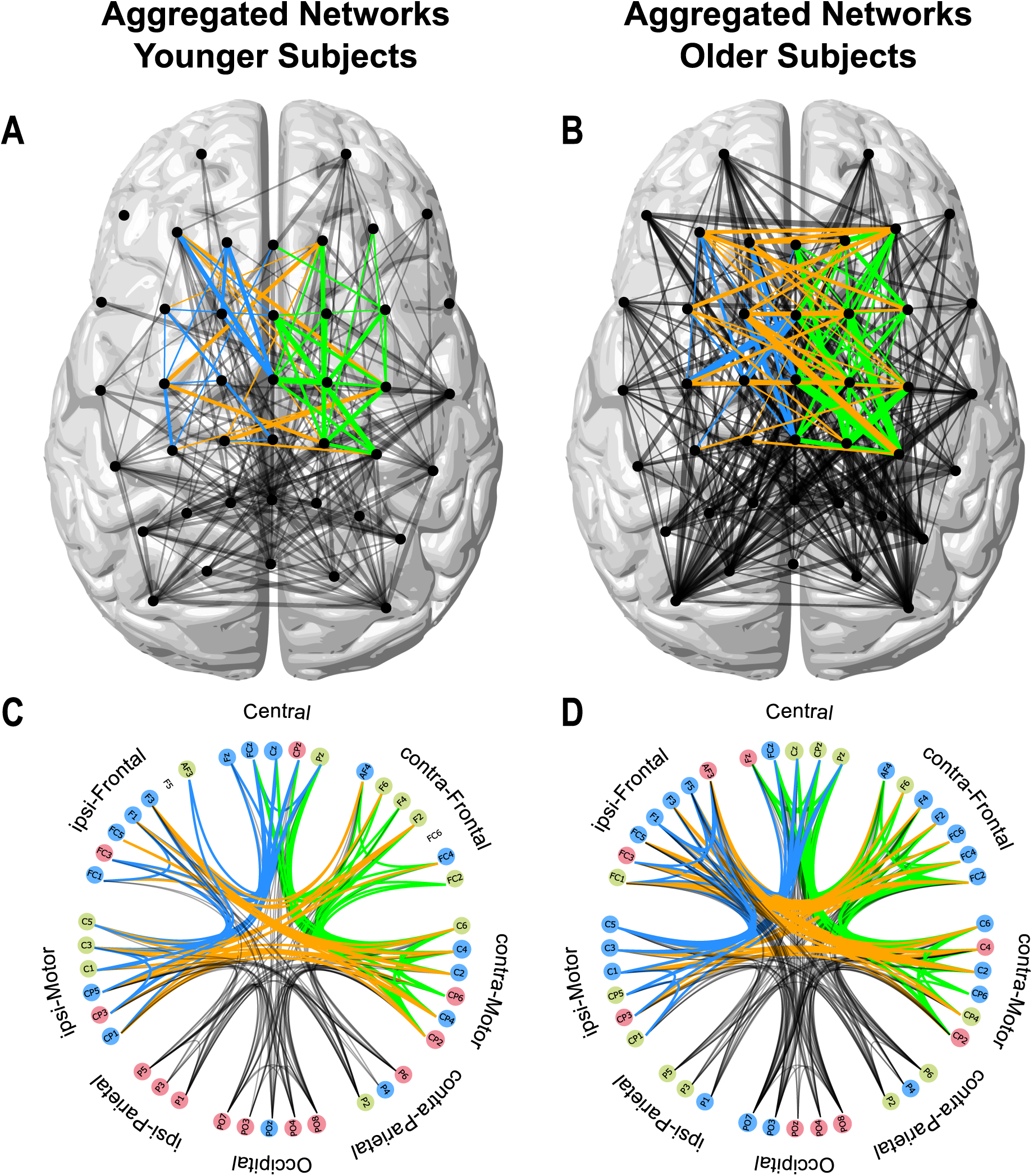
Aggregated networks in aging. Top: Representation of the aggregated networks for younger (A) and older subjects (B). The thickness of the lines accounts for the number of occurrences of the corresponding edge in the respective time interval. Bottom: Circular representation of the aggregated networks for younger (C) and older subjects (D). Nodes are color-coded depending on assigned clusters. Edges related to the core motor network are color-coded in blue (ipsilateral), green (contralateral), and orange (interhemispheric).

Additionally, we investigated the Louvain clustering for the aggregated networks (Fig. 8 C and D). Both aggregated networks could be clustered into three communities, which are represented by different colored nodes (blue, red, and green). The YS network clustered into an occipital-parietal cluster (red nodes), a cluster mainly involving the nodes above the ipsilateral frontal, central, and contralateral motor cortex (blue nodes) and a contralateral frontal and ispilateral motor regions one (green), whereas the OS network showed a cluster that mainly involved nodes above frontal areas and ipsi- and contralateral motor cortex (blue nodes), a cluster above parietal and central areas (green nodes), and one cluster containing the remaining nodes (red). Thereby, the clustering results of the aggregated networks supported the network differences of the dynamic networks during movement execution described above.

### Network information flow

In the last step, we analyzed the HUB nodes of the dynamic graphs using the betweenness centrality. We investigated the order of first appearance as well as the timecourse of the HUB nodes (Fig. 9). In both groups, occipital nodes appeared as the first HUB nodes in the system. In YS, HUB nodes were then shifted to parieto-central (light blue) and later to contralateral motor nodes. After most participants pressed the button (cf. Fig. 9 B (boxplot)), the HUB nodes switched back and forth between central/fronto-central (dark blue) and contralateral motor nodes. Whereas OS HUB nodes first shifted to central and fronto-central nodes, from there to contralateral frontal nodes, and only in the last step to contralateral motor nodes. A comparison of the timecourses showed that OS HUBs remained roughly 100 ms longer in occipital nodes than in YS. Thus, YS HUB nodes shifted about 100 ms earlier to parieto-central nodes and finally also to contralateral motor nodes.

**Fig 9.**
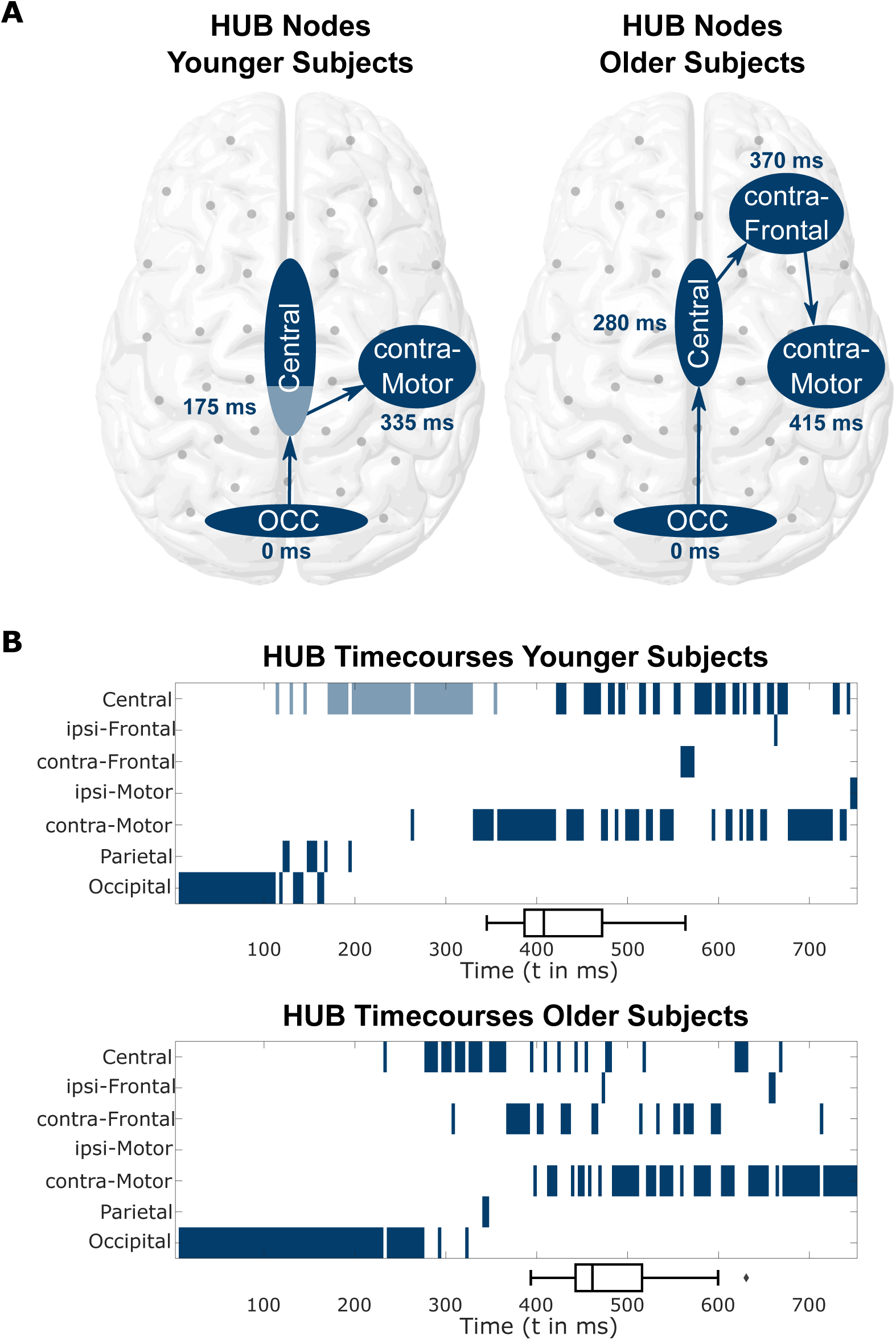
HUB nodes. Timepoints of first occurrences of HUB nodes (A) and timecourses of HUB nodes in the interval [0, 750] ms (B) - represented by groups of electrodes (Occipital: PO7, PO8; Central: Pz (light blue), CPz, Cz, FCz; contra-Frontal: FC2, FC4, FC6, F2, F4, F6; contra-Motor: C2, C4, C6, CP2, CP4, CP6) - as defined by betweenness centrality in the networks of younger and older subjects. The mean reaction times for each age group are depicted by boxplots on the x-axis.

Because of the results presented in Fig. 9, we hypothesized that the *timing of connectivity* between electrodes above contralateral motor and central regions might be a crucial factor in establishing the differences in behavioral output observed between OS and YS. For this, we investigated whether the peak time of the rPLV was related to the timing of movement output, i.e., the reaction times. Since rPLV was strongest either between C4 and Cz/FCz or CP4 and Cz/FCz, we decided to create a virtual connection by averaging over all four possible combinations. The reaction time was significantly correlated to the peak time of this virtual contralateral motor-central connection, *p <* 0.01, *ρ* = 0.42 (see Fig. 10) confirming our hypothesis.

**Fig 10.**
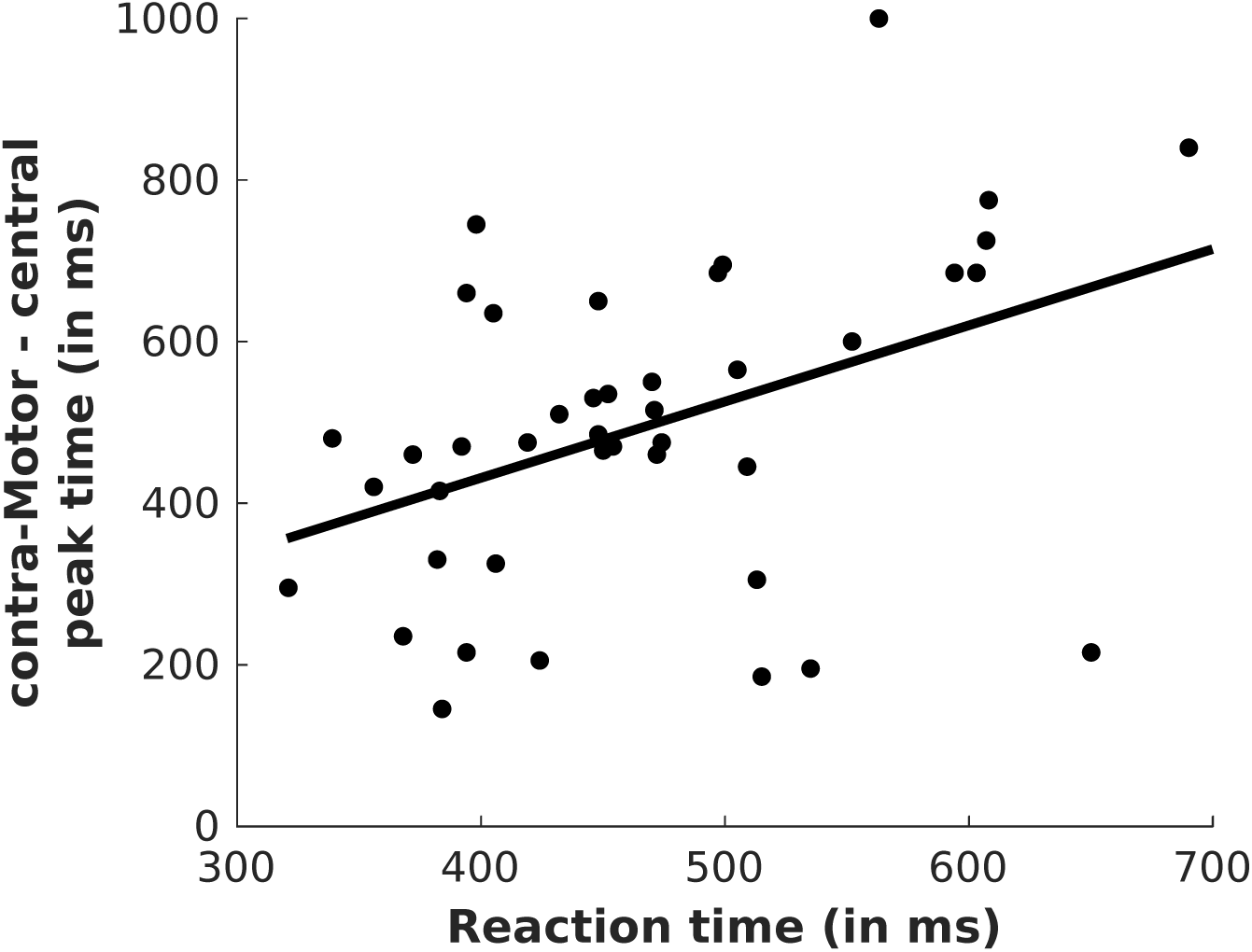
Behavioral correlation. Each subject’s reaction time (both age groups) plotted against the rPLV peak time between virtual contralateral motor and Cz / FCz electrodes.

## Discussion

In the current study we were interested in age-related changes in the processing of neural information during externally-triggered, i.e. visually-cued movements. To this extent, we performed a dynamic graph analysis on EEG data recorded from healthy younger and older participants, while they performed a visually-cued finger-tapping task. The graphs were based on phase-locking as a measure of inter-regional synchronization, which, according to the communication through coherence (CTC) hypothesis, reflects increased functional connectivity between neural populations expressed by coherently coordinated firing patterns (Fries, 2005, 2015). For this purpose, we calculated the rPLV, which measures the instantaneous synchronization between two distinct recording sites. The great advantage of rPLV is that it is capable of measuring fast transient synchronizations and thus preserves the advantage of the high temporal resolution of EEG recordings, which is essential for the construction of dynamic graphs related to the motor task. In contrast, other frequently used methods like dynamic causal modeling (DCM) or Granger causality require statistically stationary signals and thus result in connectivity information with lower time resolution. Since some of the effects we observed (e.g., decreased node flexibility) lasted for only a few hundred milliseconds and, hence, could not guarantee stationary signals (as they took place close to the button press), the application of rPLV instead of DCM or Granger causality was crucial for investigating the underlying dynamic neural networks in our study.

To start with, we computed the relative power changes of all considered electrodes in the *δ* − *θ, α* and *β* frequency range and the rPLVs between all pairs of the considered electrodes in these frequencies. Note that only rPLVs in the *δ* − *θ* frequency range turned out to be significantly increased. The electrodes were then defined as the nodes of the dynamic graph, while the edges for each time-point represented significantly increased rPLVs. We compared the time evolution of the dynamic networks that we obtained for the younger and older subjects with a focus on stimulus processing, movement preparation and movement execution. We investigated the networks considering three different aspects: the overall connectivity, i.e., the node degree of the whole graph in general and the one of the motor-related nodes, the formation of clusters, i.e., subnetworks during the different phases of the motor act, and the evolution of HUB nodes over time, i.e. the network information flow.

### Relative amplitude changes in oscillations

The analysis of the changes in event-related amplitude differences has revealed varying effects for the individual frequency bands. In the low frequencies (*δ* − *θ*), neither in the visually-cued motor nor in the contrast condition significant differences in the strength of the amplitude changes between both groups could be seen. In the higher frequencies (*α* and *β*), well known motor effects, the so-called ERD and ERS could be observed in the oscillations (Gerloff, Richard, et al., 1998; Gerloff, Uenishi, Kunieda, Hallett, & et al., 1998; Pfurtscheller & da Silva, 1999; Neuper, Woertz, & Pfurtscheller, 2006). The ERD during motor execution (approximately 500-700 ms after stimulus) differs in the mid frequency range (*α*). Here, the OS show a significantly stronger and longer ERD. However, this change was much less pronounced in the contrast as compared to the visually-cued motor condition. In the higher frequencies (*β*), a significant difference between YS and OS could be seen especially in the time after motor execution (about 1000 ms) in ERS. The YS show a clear *β*-rebound after movement execution, which is not present in OS. In contrast to the results in the *α* frequency range, this effect is visible in the visually-cued motor as well as in the contrast condition. Analogous results for aging differences in the respective frequencies, in particular in ERD and ERS have been reported (Labyt et al., 2003; Yordonova, Kolec, Hohnsbein, & Falkenstein, 2004; Toledo, Manzano, Barela, & Kohn, 2016; Liu et al., 2017). Finally, differences in movement time were about 6 ms, whereas differences in ERD lasted about 300-500 ms after movement. Thus, the ERD outlasted the end of the movement longer in OS than in YS. Hence, the longer ERD in OS cannot be explained by differences in movement time.

### Setup of dynamic networks

In the construction of the dynamic networks in the *δ* − *θ* frequency range, we defined a connection at a given time point to be present if the respective rPLV was significantly increased compared to baseline (i.e., before the stimulus) levels. Since this involved extensive testing, we applied FDR correction (q = 0.05) to account for the considerable number of multiple comparisons. One could argue that this method is not strict enough. However, it has turned out that even with a stricter FDR correction (q = 0.01), the main differences in motor-related networks between YS and OS prevailed and the core message of our results remained unaffected (results not shown). Furthermore, we decided to analyze our data at the electrode level. At least in principle, a source localization could have provided further insights into the origin of the effects reported here. However, we refrained from doing so to avoid the ill-posed inverse problem, which only provides an assumption of the source activity and could, therefore, lead to erroneous interpretations (Bradley, Yao, Dewald, & Richter, 2016; Grech et al., 2008; Wendel et al., 2009). Instead, we used the small Laplacian reference to substantially reduce the effect of volume conduction on our data. Although not a source localization method, through the small Laplacian, electrodes that are directly above the sources are maximally sensitive to the underlying activity. We have to keep in mind that thereby, the small Laplacian also eliminates possible physiological long-range connections particularly in the low frequencies. However, this means that we only underestimate the number of long-range connections and that the presented ones are likely to be of true physiological origin. Furthermore, we can see in our results, that there are many neighboring electrode pairs that do not have significant rPLVs, i.e. connections, which one would expect to see in the case of pure volume conduction (e.g. C1-Cz, C4-C6). These observations suggest that our results can also not primarily be driven by volume conduction.

### Dynamic graph connection density

Several fMRI studies have reported an overactivation related to the aging human brain (Sailer et al., 2000; Heuninckx et al., 2005; Ward et al., 2008; Dennis & Cabeza, 2011). The results of the dynamic graphs in the *δ* − *θ* frequency band reported here, i.e. the increase in node degree, might reflect *over-connectivity*, which may relate to the overactivation previously reported in fMRI studies. One might argue that this over-connectivity in OS is driven by increases in amplitude in the respective frequency band. This, however, can be ruled out due to the fact that no difference in relative change in *δ* − *θ* amplitude could be observed between YS and OS in neither the motor and nor the contrast condition. Furthermore, significant differences in relative change in *α* and *β* amplitudes occurred only after movement.

### Community structure

The analysis of the dynamic and aggregated networks revealed a substantial increase in the number of interhemispheric connections in OS. The cluster analysis of aggregated networks in YS unraveled three clusters, a cluster mainly consisting of parietal and occipital electrodes, a cluster of ipsilateral frontal, central, and contralateral motor electrodes, and a cluster that included ipsilateral motor and contralateral frontal electrodes. In OS, on the other hand, there was no clear separation into clusters possible. This might be related to the increased number of interhemispheric connections and thus a weaker inter-regional separability, which is consistent with the HAROLD model stating reduced hemispheric asymmetry during cognitive tasks (Cabeza, 2002).

Additionally, we performed a similar cluster analysis on the dynamic graphs for each time-point. The clusters were different in both groups for almost the whole analyzed period. The dynamic graph communities in YS during motor execution were, similar to the aggregated ones described above, separated in a more structured way, i.e., a dominant cluster including ipsilateral frontal, central, and contralateral motor electrodes. The most striking difference between YS and OS was observed during movement preparation, i.e., immediately before movement onset. While YS showed a peak in VI between time-points in this epoch, i.e., a very variable network structure, OS networks exhibited a drop in VI accompanied with a significant decrease in node flexibility. The decreased flexibility in OS networks might be related to the *over-connectivity* described above and the fact that OS expressed difficulties in recruiting task-specific subnetworks (Mitrushina & Satz, 1991; Babcock et al., 1997; Cabeza, 2002) and needed the whole network, i.e., higher effort to keep nearly the same task performance as YS.

### Network information flow

We furthermore included an analysis of HUB nodes of the dynamic graphs and focused on the time points of their first occurrence and their timecourses. In YS, the networks involved a direct shift of the HUB nodes from occipital via parieto-central to contralateral sensorimotor electrodes. Once YS had executed the movement, HUB nodes switched back and forth between central and contralateral motor electrodes, which might be related to the incoming sensory feedback from the button press. The HUBs of OS, in contrast, remained roughly 100ms longer in the occipital regions, then shifted to central/fronto-central electrodes and then targeted an additional HUB node above contralateral frontal electrodes, which appeared at 370 ms, i.e., approximately the same time as the shift to contralateral motor areas in YS. Finally, the connectivity to contralateral sensorimotor areas dominated, which occurred roughly 80 ms later than in YS. Here, we observed two things, on the one hand the HUB nodes in OS remained longer in occipital regions, which indicates already a delay in the processing of the visual stimulus, and on the other hand a shift to more frontal areas. This shift of the HUB nodes is compatible with the PASA (posterior-anterior shift in aging) phenomenon (Dennis & Cabeza, 2011), which refers to the fact that the overall activity in OS is shifted to more frontal areas representing a stronger involvement of cognitive control.

### Connecting network changes with behaviour

The motor output itself, i.e. the movement of the index finger, did change only slightly. The difference between younger and older subjects’ movement times, i.e. the times from movement onset to the button press, was about 6 ms which is approximately one sampling point at 200 Hz of the EEG signal. The reaction times between younger and older subjects differed by approximately 70 ms, thus older subjects started their movements later than younger subjects did and therefore a time period existed in which younger and older brain networks received different input in terms of sensory feedback. However, we do not expect this to have an influence on network dynamics before movement onset where the main differences in network structures were observed. The reaction time on the other hand is closely related to the processing of the stimulus. We hypothesize that additionally to the prolonged processing of the stimulus in occipital regions, a *detour* via frontal areas in particular is accountable for the longer reaction times in OS. This hypothesis is supported by the reported positive correlation between the reaction times and the peak timing of the connection between central and contralateral motor electrodes. This result suggests a decreased behavioral output in case of a stronger PASA effect. We therefore hypothesize that increased frontal activity reflects reduced efficiency or specificity rather than compensation (Morcom & Henson, 2018).

### Comparing motor network connectivity between visually-cued and *voluntary* movements

In a previous publication (Rosjat et al., 2018), we reported on an invariant, i.e. present and unchanged in all movement phases, motor network in younger volunteers consisting of iPFC/iPM, mPFC/SMA, and cSM while subjects were performing voluntary, i.e. internally triggered, movements. This invariant network was significantly attenuated in OS and accompanied by increased interhemispheric connectivity. The results we have presented here reveal comparable patterns, albeit in the execution of an externally triggered motor task (as opposed to the voluntary movements before). YS established most (motor-related) connections between ipsilateral frontal, central, and contralateral motor regions. The networks of OS, on the other hand, showed additional interhemispheric connections, especially between ipsilateral frontal and contralateral frontal as well as ipsilateral motor and contralateral motor electrodes. Also, similar to the results during voluntary movements, OS exhibited higher connectivity between central and contralateral frontal electrodes. These results indicate that by investigating the contrast between visually-cued and vision-only conditions, we devised a condition alike the pure motor task (voluntary movement). When contrasting visually-cued and vision-only conditions, it cannot be excluded that NOGO effects, i.e. effects of actively suppressing a prepared movement, may occur in the vision-only condition. However, the fact that false movements were observed only in a negligible number of cases in both younger and older subjects indicates that the subjects did not prepare for any movement in the vision-only condition. If they had done so, both conditions would have been comparable or even identical up to the time of movement initiation, which would have resulted in similar phase-locking, i.e. networks consisting of only a small number of connections. Since this was not the case, our data suggests that a NOGO effect played a minor role at most in our study. Thus, our findings indicate that we identified a highly stable core network (iPFC/iPM - mPFC/SMA - cSM) that is important for movement execution over several experimental conditions.

### Future perspectives

There are several possible ways to continue to further deepen our knowledge on aging-associated functional changes in brain networks underlying movements. One might include a group with pathological aging (e.g. stroke) to investigate the compensation following brain lesions. This would be particularly interesting in a longitudinal study, which may serve to investigate how the compensation mechanisms develop while motor skills improve in rehabilitation. An attempt could be made to investigate subjects with good and subjects with poor motoric improvements in rehabilitation to try to identify advantageous and disadvantegeous changes in information flow in the different movement phases. It could then be explored whether influencing these could lead to an improvement in motor response. Another interesting approach would be to include a more challenging motor task (e.g. complex movement patterns) to get a better understanding of whether a detour via frontal nodes really reflects a reduced efficiency in recruiting specified brain regions or whether it might be beneficial for older subjects.

## Summary

In this article we performed an analysis of dynamic networks based on synchronization in the *δ* − *θ* frequency range in the context of stimulus processing and motor execution in order to understand the neural mechanisms underlying HAROLD and PASA. We were able to demonstrate a similar structure as has been shown in self-initiated movements indicating an independence of the underlying neural mechanisms on movement initiation: The networks in older subjects displayed an overall increased connectivity, especially in motor related electrodes by establishing significantly more interhemispheric connections which is consistent with the HAROLD model. In addition, we could show that the flexibility in the networks of older subjects decreased, indicating a kind of over-connectivity and resulting difficulties in recruiting task specific regions. Finally, an analysis of the HUB nodes, i.e. the neural information flow revealed an extended stimulus processing in occipital regions and a detour via frontal regions in older subjects, which is compatible with the PASA phenomenon.

## Conflict of interest

The authors declare no conflict of interest.

## Data availability statement

Data has been made available at https://doi.org/10.26165/JUELICH-DATA/T1PKNZ. The specific codes used for automated computation of dynamic graphs and the corresponding measures can be shared upon reasonable request.

## Acknowledgements

This study was funded by the University of Cologne Emerging Groups Initiative (CONNECT) within the framework of the Institutional Strategy of the University of Cologne and the German Excellence Initiative. SD gratefully acknowledges additional support from the German Research Foundation, Germany (DA1953/5-2). GRF gratefully acknowledges additional support from the Marga and Walter Boll Foundation.

## Notes

### Competing Interest Statement

The authors have declared no competing interest.

