## Supporting Video 1 for "Stimulus transformation into motor action: dynamic graph analysis reveals a posterior-to-anterior shift in brain network communication of older subjects"

**Supporting Information**

We performed a dynamic graph analysis in the interval from 0ms to 1000ms in younger and older subjects. In the main manuscript we present snapshots of the dynamic graphs (Fig. 5) and the aggregated networks (Fig. 8). The Supplementary Video V1 displays the whole timecourse of the dynamic graphs for younger and older subjects

Supplementary Video V1

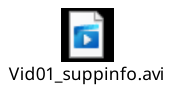

*Full timecourse of the dynamic graphs resulting from the rPLV analysis for younger subjects (left) and older subjects (right). Motor related connections are presented in blue (ipsilateral), green (contralateral) and red (interhemispheric).*
